# Sunk cost sensitivity in mice, rats, and humans on the Restaurant Row and WebSurf tasks cannot be explained by attrition biases alone

**DOI:** 10.1101/2021.10.07.462802

**Authors:** A. David Redish, Brian M. Sweis, Samantha Abram, Anneke Duin, Rebecca Kazinka, Adrina Kocharian, Angus MacDonald, Brandy Schmidt, Neil Schmitzer-Tobert, Mark Thomas

**Affiliations:** Department of Neuroscience, University of Minnesota; Nash Family Department of Neuroscience, Friedman Brain Institute, Icahn School of Medicine at Mount Sinai, New York, NY 10029; Department of Psychiatry, Icahn School of Medicine at Mount Sinai, New York, NY 10029; San Francisco Veterans Affairs Medical Center, 4150 Clement Street, San Francisco, CA 94121; Department of Psychiatry and Behavioral Sciences, University of California San Francisco, 505 Parnassus Avenue, San Francisco, CA 94143; National Institutes of Health, 9000 Rockville Pike, Bethesda, Maryland 20892; Graduate Program in Clinical Science & Psychopathology Research, University of Minnesota; Graduate Program in Neuroscience, University of Minnesota, and Medical Scientist Training Program, University of Minnesota; Department of Psychology, University of Minnesota; Department of Psychology, Wabash College

## Abstract

In a recent bioRxiv preprint, Ott et al. argue that sensitivities to sunk costs that have been reported in two serial foraging tasks (the Restaurant Row task in mice and rats, and the Web-Surf task in humans) may be due to simple consequences of the way that subjects perform these tasks and not due to an actual sensitivity to sunk costs. However, several variants of these tasks have been studied, in which the sensitivity to sunk costs changes. In order to test the Ott et al. model against these experimental observations, we simulated the model under these additional experimental conditions. We find that it is incompatible with the actual data. While we applaud the simplicity of the Ott et al. model, we must reject it as an explanation for the observed sensitivity to sunk costs seen in these tasks. We thus conclude that the alternative explanation - that mice, rats, and humans are sensitive to actual sunk costs in these tasks - is a better explanation for the data.

## Background

### Restaurant Row

In 2014, we introduced a new foraging task in which rats experienced a sequence of delay-to-reward offers and made *accept / skip* decisions for each offer. Because the rats had a limited time to gather their complete food complement for the day, time was a limited resource, making this an economic task. We found that rats showed regret-like behaviors in which they responded differently to losses due to mistakes of their own agency (regret) vs losses that arose from environmental conditions (disappointment) (Steiner and Redish 2014). In this task, rats ran around a cyclical (octagonal) environment, encountering rewards at each of four different “restaurants”, each providing a different flavor of reward. As the animal entered a restaurant, a series of tones played (1/sec) such that the pitch indicated the delay to the reward and the descending pitches counted down to the reward. Rats could wait out the delay or leave before the countdown completed. If the rat left the restaurant before the delay completed, the offer was rescinded and the rat had to proceed to the next restaurant to receive another offer. At each restaurant, rats demonstrated an individual delay threshold, below which they accepted offers by waiting out the delay, and above which they skipped by leaving the offer zone relatively quickly. These thresholds were different for each rat for each restaurant, indicating that the decisions were based on an interaction of cost and flavor, but they were also generally consistent across days for a given rat for a given restaurant, indicating an economically-consistent revealed preference. These basic results have been replicated numerous times in both rats and mice (Schmidt, Duin, and Redish 2019; Sweis, Larson, et al. 2018; Sweis, Redish, and Thomas 2018; Sweis, Thomas, and Redish 2018; Sweis, Abram, et al. 2018; Schmidt and Redish 2021).

### Web-Surf

We then developed an analogous task for humans in which humans foraged for rewarding videos with four “galleries” providing a different video category that would only play after a delay bar completed as if downloading content on the web (Abram et al. 2016). We found that humans showed similar behaviors, including individual preferences, indicated by decision thresholds that depended on an interaction between delay and video category. We also asked humans to rate the videos after watching them and to rank the categories at the end of the session. All three economic preference measurements (decision thresholds, video ratings, category rankings) were correlated with each other, indicating that the task was measuring economically consistent preferences. Again, these results have been replicated numerous times, both using in-person and in online versions (Sweis, Abram, et al. 2018; Abram et al. 2019; Haynos et al. 2019; Huynh et al. 2021; Kazinka, MacDonald, and Redish 2021). Similar results have been found with still photo rewards (Abram et al. 2016) and with food rewards (Huynh et al. 2021).

### Two-phase variant with separate offer and waiting zones/phases

We then developed an important variant of these two tasks in which there were separate offer and wait zones (Sweis, Thomas, and Redish 2018). On entering the offer zone, the subject was informed of the delay (by a repeating tone of a given pitch), but the countdown did not decrease while subjects remained in the offer zone. Subjects could skip as before, leaving the offer zone for the next restaurant / gallery or they could accept by entering the wait zone, at which point the delay began to count down (as in the original Restaurant Row task). Importantly, subjects in this variant could quit out of the wait zone even though they had accepted the offer by entering into the wait zone, at which point the countdown stopped and the offer was rescinded, as if they had decided to skip. We have run several cohorts of this two-phase variant on rats and mice (Sweis, Thomas, and Redish 2018; Sweis, Abram, et al. 2018; Sweis, Redish, and Thomas 2018)

We created an equivalent version in Web-Surf, in which subjects had two phases: an offer phase and a wait phase. In the offer phase, subjects were informed of the delay by a download bar and a number, but the download bar did not start decreasing until the subjects accepted the deal and entered the wait phase. Importantly, subjects could again, quit out of the wait phase. We have run several cohorts of humans on this two-phase variant both in-person and online (Sweis, Abram, et al. 2018; Kazinka, MacDonald, and Redish 2021; Huynh et al. 2021). We will use the terms “offer zone” / “offer phase” and “wait zone” / “wait phase” interchangeably in this document.

### Sunk Costs

Analyzing behavior on this two-phase variant, we found that the decision to *quit* depended not only on the time remaining in the countdown but also on the time spent in the wait phase - that the longer the subject had spent within the wait phase, the less willing they were to quit, even for identical future conditions (Sweis, Abram, et al. 2018; Kazinka, MacDonald, and Redish 2021). Time already spent waiting is the definition of <u>sunk costs</u> (Staw 1976; Staw and Fox 1977; Staw and Ross 1989). A <u>sensitivity to sunk costs</u> is commonly identified as an economic fallacy --- theoretically, decisions should be based on the future conditions. Nevertheless, we found that time already spent (sunk costs) affected the decisions, leading to escalation of commitment (increased willingness to continue a course of action). An escalation of commitment occurs when the longer the agent has spent in the situation, the more committed the agent is to the situation (the less likely the agent is to quit), see (Staw 1976). A sensitivity to sunk costs (increasing costs sunk leads to decreases in quitting) is a form of escalation of commitment.

In a recent bioRxiv preprint, Ott, Masset, Gouvêa, and Kepecs proposed that this observed behavioral sensitivity to sunk costs could be an epiphenomenon consequence of a combination of attrition statistics, a drift diffusion model, and a changing threshold for quitting (Ott et al. 2021). In this paper, we analyze their model and compare it to the extensive data we have from Restaurant Row, Web-Surf, and follow-up variants (such as variants with and without an offer phase (Steiner and Redish 2014; Schmidt, Duin, and Redish 2019; Sweis, Abram, et al. 2018), the Movie Row task, in which subjects navigate spatially in virtual reality to view movies (Huynh et al. 2021), the Candy Row task, in which subjects wait out delays to receive small pieces of candy (Huynh et al. 2021), and the Known-Delay/Randomized-Delay tasks, in which we manipulated the predictability of the upcoming delay in each restaurant (Duin et al. 2021)). While we applaud the mechanistic simplicity of the (Ott et al. 2021) model, we find that their model does not fit key aspects of the data. We therefore conclude that their model does not explain the sunk cost behavior that we are observing in these tasks.

## Their model

(Ott et al. 2021) propose that they can replicate the sunk cost behavior reported in (Sweis, Abram, et al. 2018) with a simple model in which they assume (1) that the subject has a *willingness to wait* ***W*** which distributes normally (W = W_0_ + N(σ_W_)), (2) that the decision to enter an offer occurs when ***W*** is greater than the threshold for that offer ***T***_OZ_, (3) that ***W*** then wanders with a given variance (σ_N_), and the subject quits if ***W*** ever crosses a quit-threshold in the wait zone ***T***_WZ_. Although they do not discuss the importance of the shape of the quit threshold ***T***_WZ_ in their preprint, they assume that the quit-threshold ***T***_WZ_ decreases with time so that it starts at the entry threshold ***T***_WZ_(t=0) = ***T***_OZ_ and reaches 0 at the time of reward ***T***_WZ_(t=***T***_OZ_) = 0. We find that this decreasing quit-threshold drives a large part of the sunk cost behavior in their model.

We have built a simulation of their model and agree that it shows the basic effect of sunk costs, replicating the basic effect that the slope of the probability of earning a reward as a function of the time-remaining decreases as a function of the time already waited. For each accepted offer, we measure the probability that the agent waits out the delay at each second. This produces a probability of earning the reward, or **p(Earn)** for every combination of time-remaining and time-spent. We plot these in several formats, as shown below.

We concur that their model does show a sensitivity to sunk costs. However, two important questions remain: First, *what are the factors within their model that create this sensitivity to sunk costs?* Second, *how well does their model describe the data we have observed in mice, rats, and humans?*

They claim that the sensitivity to sunk costs arises from the *<u>selection bias</u>* inherent in their model (which they claim is a function of the task design) - accepting an offer with a high threshold implies that ***W*** was selected from the higher part of that distribution, which implies that there is attrition in the distribution of ***W*** on accepting an offer. However, our examination of their model under different conditions finds that their model continues to show sunk costs, even when the agent takes every choice. Thus, there must be other factors in their model producing sunk cost sensitivity even without the attrition bias.

The results shown in Figure 5 imply that the attrition hypothesis (that accepting a higher offer implies a higher ***W*** on acceptance of the offer and entry into the wait zone) is not the primary factor driving sunk costs in their model. Thus, we set out to look for what else was producing a sensitivity to sunk costs in their model.

The other factor that is critical to the presence of sunk costs in their model is that their model includes a changing **quit threshold *T***_**WZ**_. This quit threshold as shown in their Figure 2 decreases from being equal to the offer on entry into the wait phase to 0 at the time of reward.

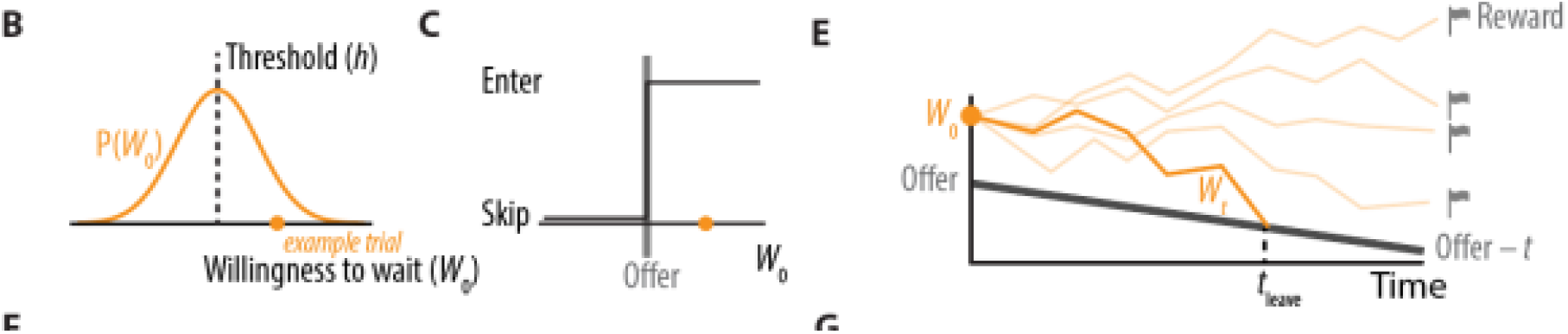
Figure reproduced from Ott et al. Their Figure 2 showing the key components of their model.

**FIGURE 1:**
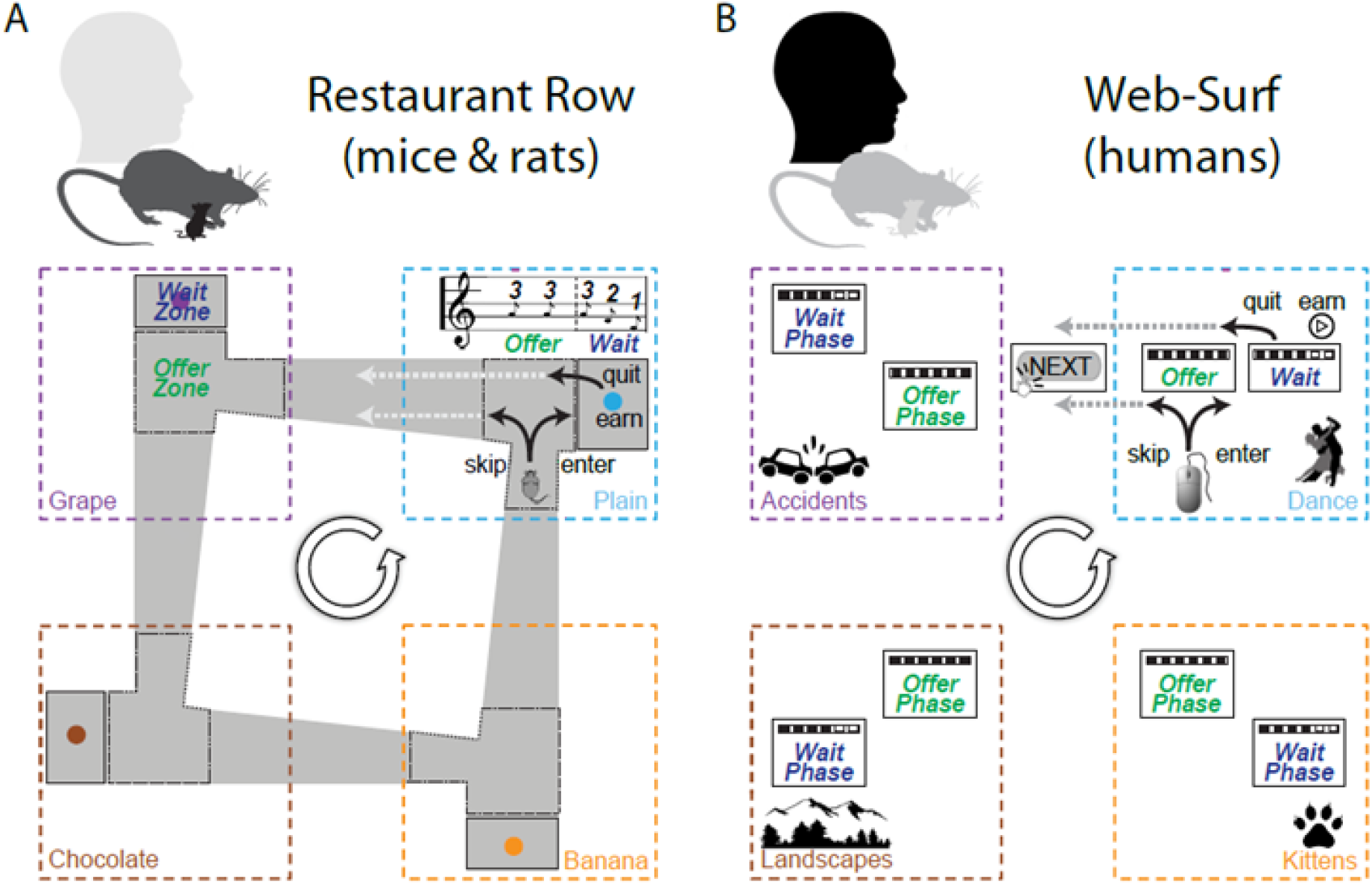
Restaurant Row and Web-Surf. A: The Restaurant Row task for mice and rats. As the animal enters the offer zone, the delay is indicated by a tone, which only starts counting down on entering the wait zone. Animals encounter restaurants serially by proceeding around the cycle counterclockwise. B: The Web-Surf task for humans. An offer is provided to the human as a download bar with a set delay, but the download does not start to count down until they select “enter” to enter the wait phase. Humans encounter the video galleries serially by clicking through a sequence of buttons. Note the topological analogies. Figure from (Sweis, Abram, et al. 2018). Used with permission of the publisher.

**FIGURE 2:**
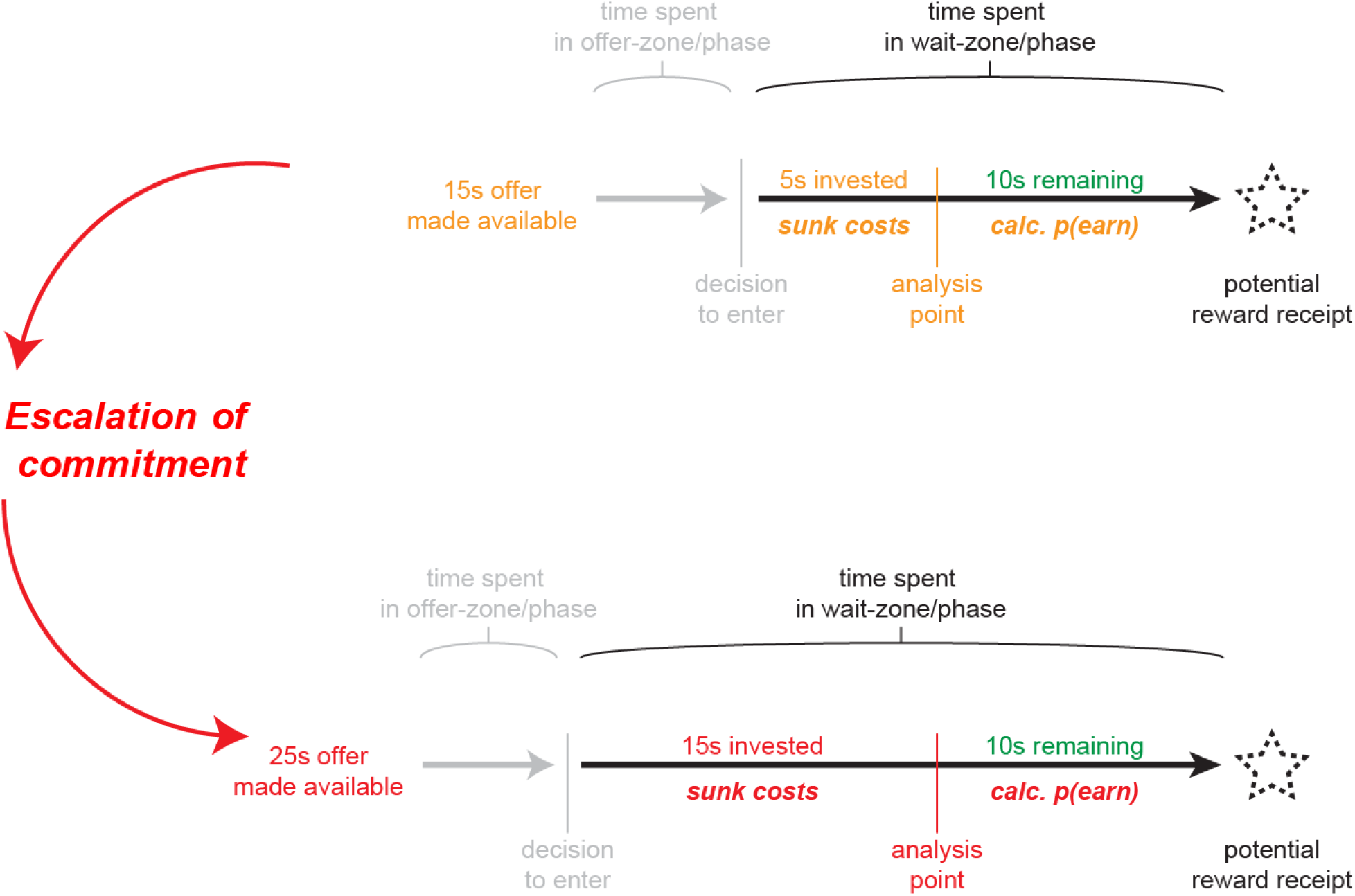
Conceptualizing sunk costs on the Restaurant Row and Web-Surf tasks. At a given point within the countdown in the wait phase, we ask what is the likelihood that the subject will wait out the delay to receive the reward. This is ***p(earn)***. In the two examples, the future from the two analysis points is identical (10s remaining before receiving the same reward), however, the time spent in the wait zone (the sunk costs) is different. If we find a difference in the willingness to wait out the future delay, *p(earn)*, then we can conclude that the decision has included some sensitivity to the sunk costs. Figure from (Sweis, Abram, et al. 2018). Used with permission of the publisher.

**FIGURE 3:**
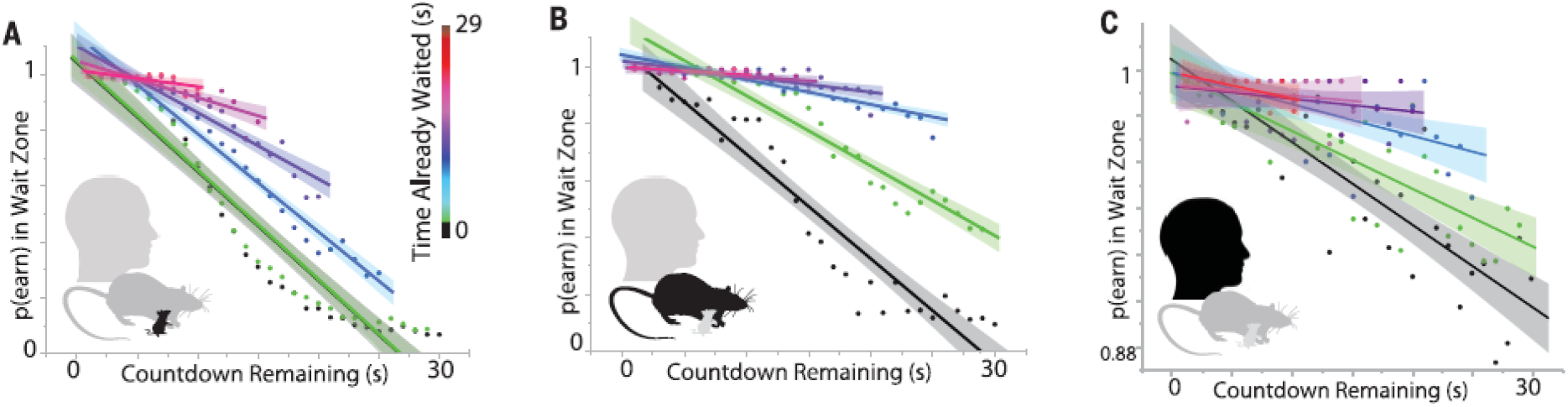
Observation of sunk costs within all three species. On each plot, the graph shows the probability of earning, *p(earn)*, as a function of the time remaining in the wait zone. Each colored dot and each line aligned to the colored dots corresponds to the p(earn) given that the subject has waited a certain amount of time in the wait zone already. Notice that the slopes of the lines decrease, indicating that having waited longer, subjects are more likely to wait out the delay, even for a given countdown remaining. Thus, following the logic in Figure 2, we conclude that the subjects are behaviorally sensitive to sunk costs. Figure from (Sweis, Abram, et al. 2018). Used with permission of the publisher.

**FIGURE 4:**
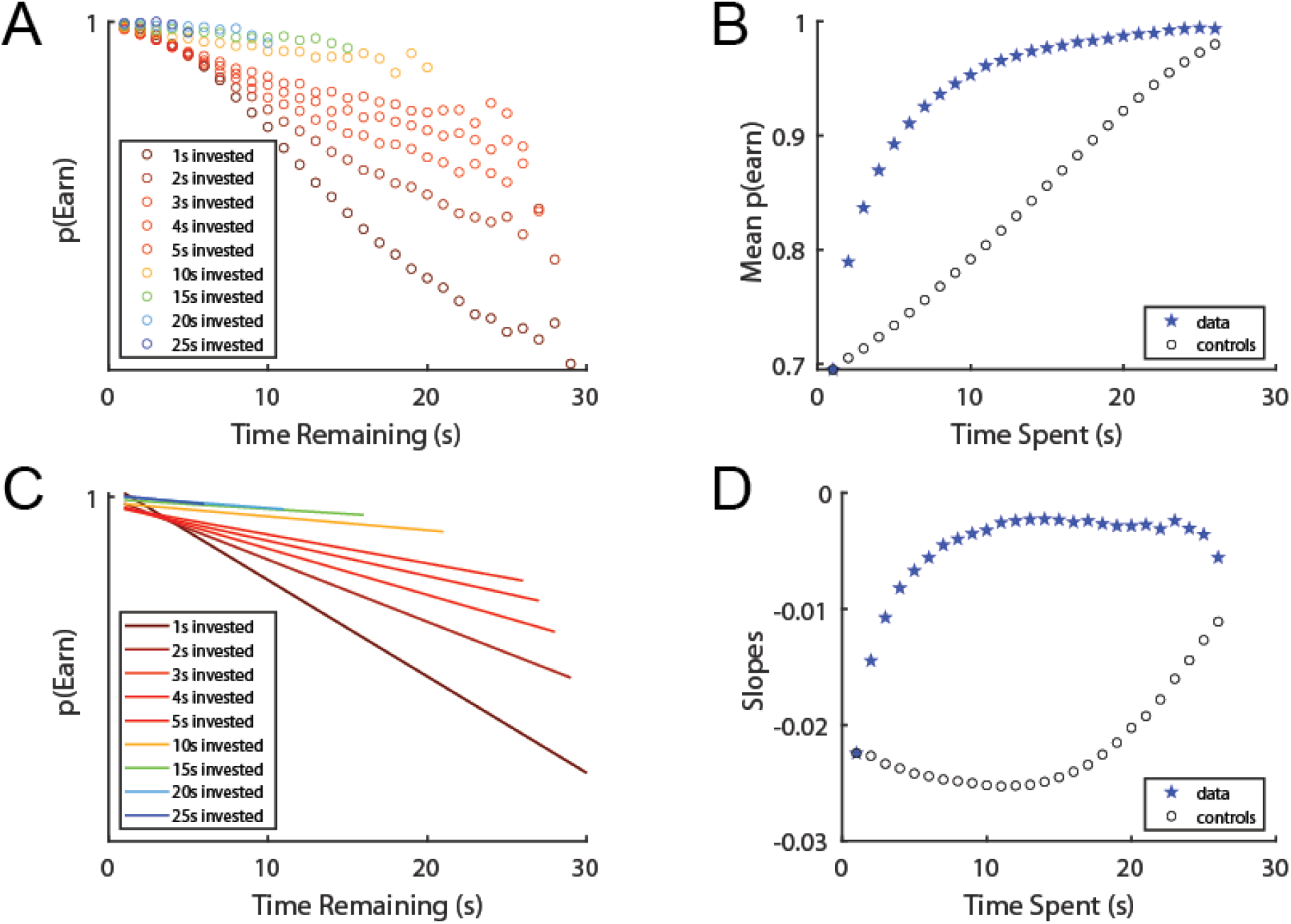
Reproducing the basic structure of the (Ott et al. 2021) model. (A) The ***p(Earn)*** matrix. Each dot shows the probability of staying through the delay as a function of time remaining (x-axis) and time-spent (color). (B) Taking the mean across each time-spent and the equivalent set of points with 0s invested shows increased **p(Earn)** as a function of time invested. (C) Alternatively, we can measure the linear slope of each time invested, revealing that the slopes flatten with time-invested. This is how data was displayed in (Sweis, Abram, et al. 2018). (D) Comparing the slope with the slope of the equivalent set of points with 0s invested shows decreased slopes with increased time-invested. This is how data was displayed in (Sweis, Abram, et al. 2018). **<u>*Results shown are from our simulations using the Ott et al. model*</u>**.

**FIGURE 5:**
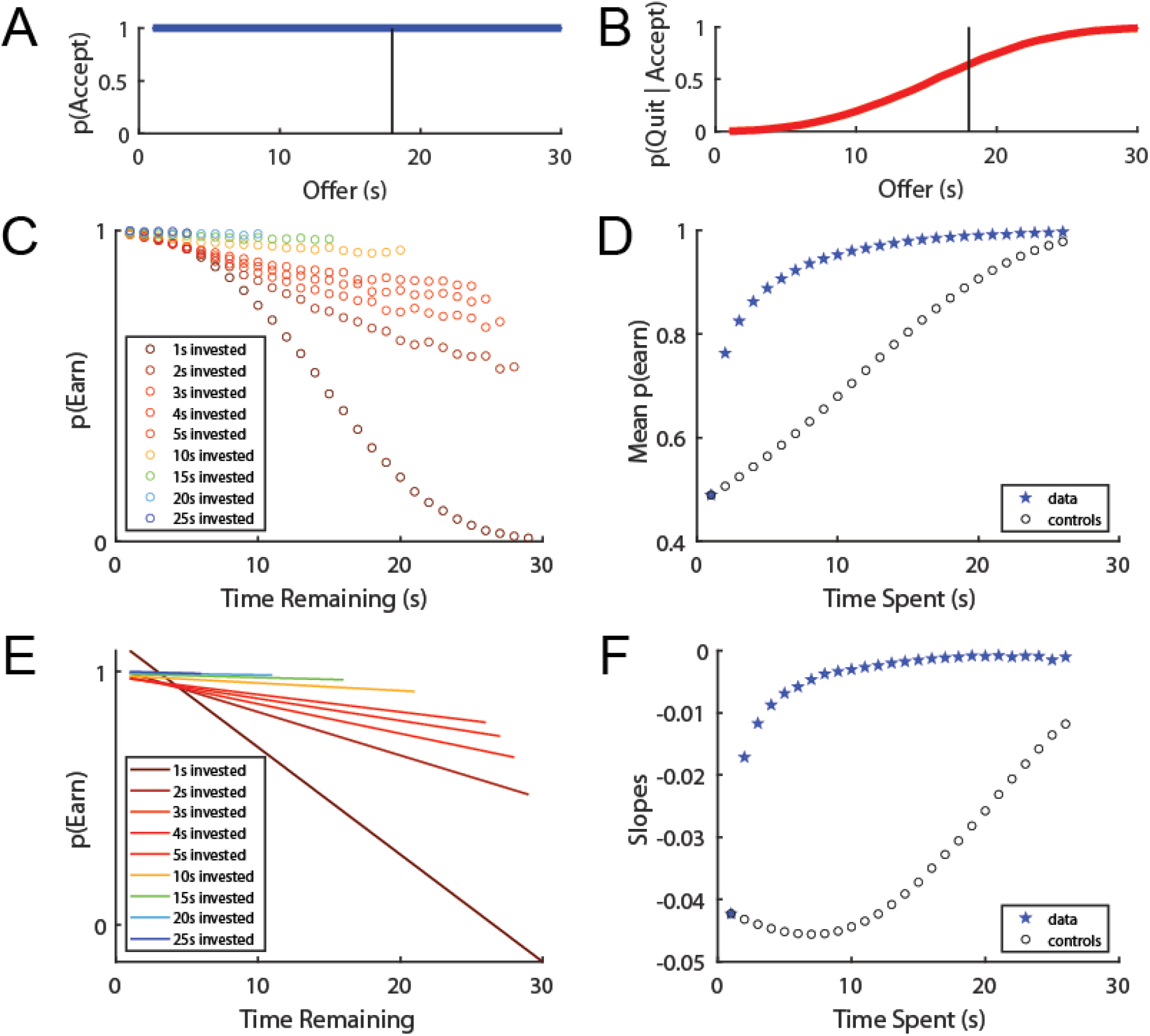
Simulation of Ott et al. model with p(accept) = 1.0. To directly address the selection bias hypothesis that Ott et al. propose, we examined the sensitivity to sunk costs in the model when the agent accepts every offer, thus removing selection bias as a factor. (A) All offers are accepted. The Ott et al. model assumes that the threshold = 18s, indicated by the black line. (B) However, the agents still show a continuous quitting function centered slightly below the threshold. (C-F) Panels as in Figure 4. The Ott et al. model continues to show a robust sensitivity to sunk cost, despite the removal of the selection bias that they hypothesize produces attrition, the ***W***_0_ distribution on acceptance of the offer and entry into the wait zone. **<u>*Results shown are from our simulations using the Ott et al. model*</u>**.

The quit threshold decreases at a constant rate of 1/s. Note that agents enter the wait zone with a willingness to wait, **W**, that is going to be above the offer (that’s the process by which their model accepts an offer). As the agent spends time in the offer zone, ***T***_***WZ***_ decreases so that it will be 0 when the countdown stops. This means that ***T***_***WZ***_ moves away from **W** at a rate of 1/s as the agent waits in the wait zone, making the agent less likely to quit with time in the wait zone.

Importantly, this means that the changing threshold contains information about <u>both the time remaining and the time spent</u>. When we remove this rate of change (removing the information about time spent as we remove the information about time remaining), we find that this decreases the sensitivity to sunk costs. If we set the quit threshold to the offer, we decrease the sensitivity to sunk costs. If we set the quit threshold to 0, the agent never quits. We can manipulate the sensitivity to sunk costs in their model by changing the **slope of the quit threshold**. A large part of the reason that their model is sensitive to sunk costs is that they directly built in such a sensitivity in the changing quit threshold.

In order to completely remove sunk costs in their model, we need to remove both the attrition factor that comes from the selection bias and also the decreasing quit threshold, at which point the simulations do not fit data we have observed in any of the task variants.

### Their model is fragile

In general, we have found that very small changes in their model parameters have dramatic effects on their results. For example, small changes in the parameters (for example changing σ_W_ or σ_N_) have large effects that produce very different behavior patterns unlike what we typically see in real subjects on these tasks. A fragile model per se does not mean it is incorrect, however, it does raise concerns about its capacity to characterize a variety of behavioral observations relating to sunk costs, and suggests that more robust models may better explain the observations.

## Comparing the Ott et al. model to data

In order to test their model against the available data, we run a series of simulations attempting to address variations of the Restaurant Row and WebSurf tasks.

### 1. The data finds no sensitivity to sunk costs in the offer phase

The first and most important result, which was originally observed upon introduction of the offer zone variant (Sweis, Abram, et al. 2018), is that we do not see a sensitivity to sunk costs in the offer zone, even though the subjects are informed of the delay in the offer zone, they spend time in the offer zone, and the time in the offer zone is included in the total economic time budget. This result has been replicated in online samples (Kazinka, MacDonald, and Redish 2021; Huynh et al. 2021).

(Ott et al. 2021) already acknowledge in their preprint that their model cannot explain behavior in the variants of Restaurant Row and Web-Surf that have a separate offer phase, and agree with our conclusions that decisions made in the offer and wait phases arise from different decision processes. As we have argued elsewhere (Sweis, Thomas, and Redish 2018; Kazinka, MacDonald, and Redish 2021; Sweis, Abram, et al. 2018), we agree that behavior in the offer and wait phases arise from different decision processes, consistent with both multiple-decision theories (Redish 2013, 2016) and commitment hypotheses (Gollwitzer, Heckhausen, and Steller 1990; Achtziger and Gollwitzer 2007). Although Ott et al. acknowledge that offer and wait phases may well be different, they do not provide any explanation as to why the two phases are different as to sunk costs. Moreover, it is not rational to spend any time in the offer zone, as any time spent making decisions in the offer zone could be more productively spent being made in the wait zone, while the delay is counting down. However, all three species spend a significant amount of decision time in the offer zone, and mice, at least, increase their time spent in the offer zone with experience (Sweis, Thomas, and Redish 2018; Sweis, Abram, et al. 2018).

### 2. The data finds that sensitivity to sunk costs are delayed in variants without an offer phase

The more parsimonious explanation is that subjects make a separate decision before committing to a choice and then show a sensitivity to sunk costs only after commitment (Sweis, Abram, et al. 2018; Sweis, Thomas, and Redish 2018; Kazinka, MacDonald, and Redish 2021; Gollwitzer, Heckhausen, and Steller 1990; Achtziger and Gollwitzer 2007). This hypothesis suggests that one should see this difference, even without an explicit offer phase. And, in fact, that is what we find.

The early versions of the Restaurant Row task did not have a separate offer zone or offer phase, and only included a wait zone/wait phase (Steiner and Redish 2014; Schmidt, Duin, and Redish 2019). A reanalysis of these data found that sunk costs did not start accruing for several seconds (Sweis, Abram, et al. 2018). We hypothesized that animals were spending the first few seconds making an *accept/skip* decision as if they were in an offer zone rather than a *quit/remain* decision. After making that *accept/skip* decision, they transitioned to a *quit/remain* decision and began building sunk costs. We found robust sunk costs in these animals, but only after a delay, consistent with typical decision times made in variants with an explicit [separate] offer phase.

In a recent variant without an offer phase, we directly manipulated the upcoming uncertainty of the future outcomes (Duin et al. 2021). In two parallel tasks, rats approached a series of reward sites which either had a set delay (different across the four reward sites, but constant within each reward site for a given day, changing from day to day) or had a random delay to reward (1s-30s random on each entry). In the first task (*Known-Delay [KD]*), rats knew what the upcoming delay would be and showed behaviors indicating that they had already made their decision before entering the reward zone - they maintained a fast speed on approach to offers they skipped. In the second task (*Random-Delay [RD]*), rats slowed down on every entry into an offer and then only sped up to leave after a few seconds of consideration. Furthermore, decisions in the KD task were more self-consistent, that is, rats were more likely to either choose to stay or skip for a given delay on the KD task. We found that sunk costs were decreased in the KD task, but were still present. Importantly, however, we also found that sunk costs did not start accruing in the RD task until after the decision-time (5s), while sunk costs started accruing in the KD task immediately.

The Ott et al. model contains information about inherent uncertainty in the variance of the ***W*** parameter. Thus, we modeled the difference between the KD and RD by decreasing the variance of the willingness-to-wait σ_W_. As can be seen in Figure 8, interestingly, this result did decrease the sunk costs (as attrition is a part of the effect that generates sunk costs in the Ott et al. model), consistent with the decreased sunk costs seen in KD relative to RD. Interestingly, setting σ_W_ to 0 did show a similar low sensitivity to sunk costs that broke down as the delays got large, similar to what was seen in KD. Of course, with σ_W_ = 0, there can be no “attrition” effect; instead, the entire sunk cost effect is carried in the Ott et al. model through the decreasing quit-threshold slope ***T***_WZ_.

**FIGURE 6:**
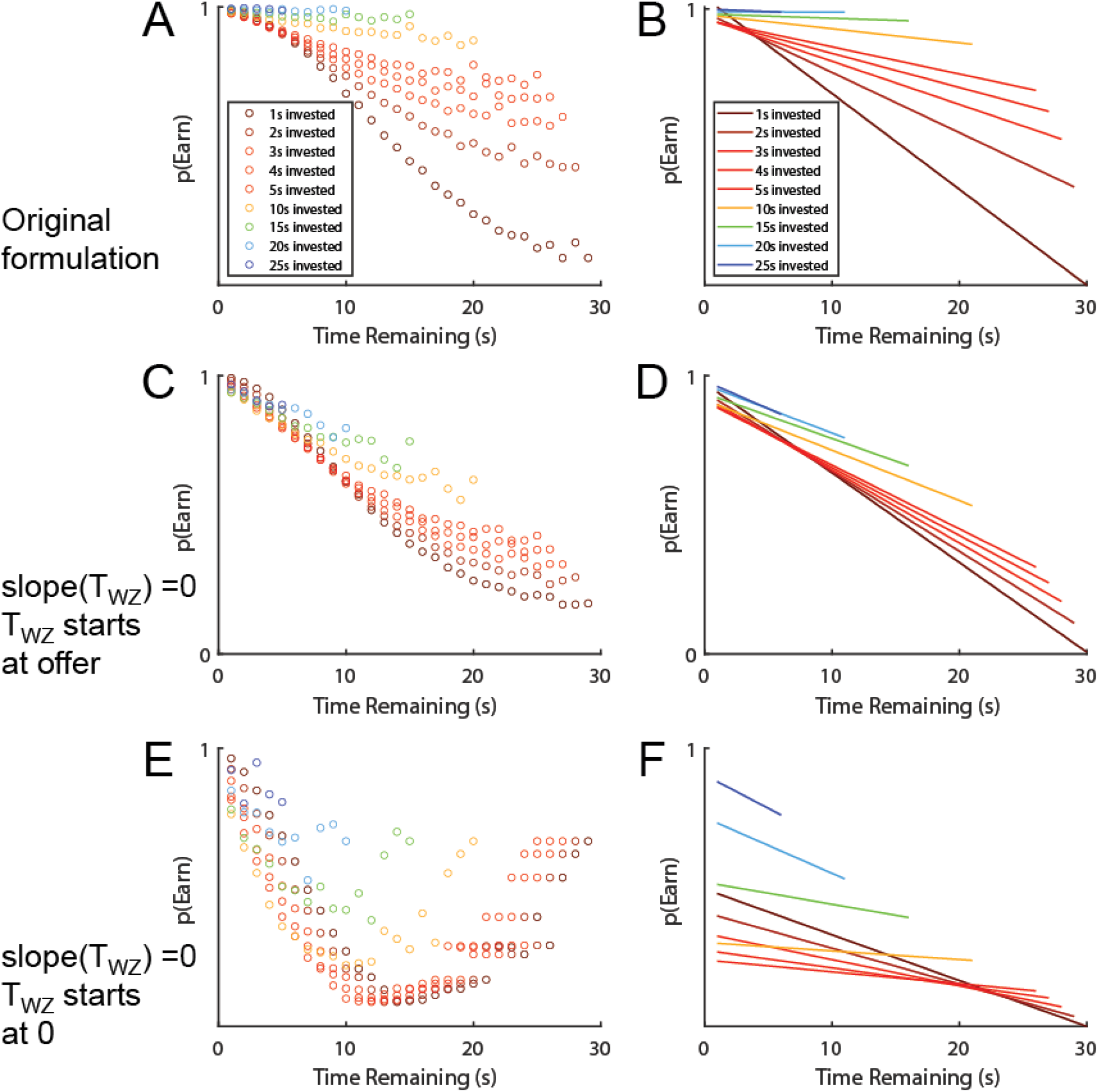
Effect of changing the slope of the quit threshold ***T***_***WZ***_ in the Ott et al. model. (A,B) the original formulation. (C,D) Effect of removing the slope of ***T***_***WZ***_, while starting ***T***_***WZ***_ at the offer, thus making the quit threshold = offer. (E,F) Effect of removing the slope of ***T***_***WZ***_, while starting ***T***_***WZ***_ to 0, thus setting the quit threshold to 0. Note that because ***W*** can wander to a number less than 0, it is still possible for the agent to quit. **<u>*Results shown are from our simulations using the Ott et al. model*</u>**.

**FIGURE 7:**
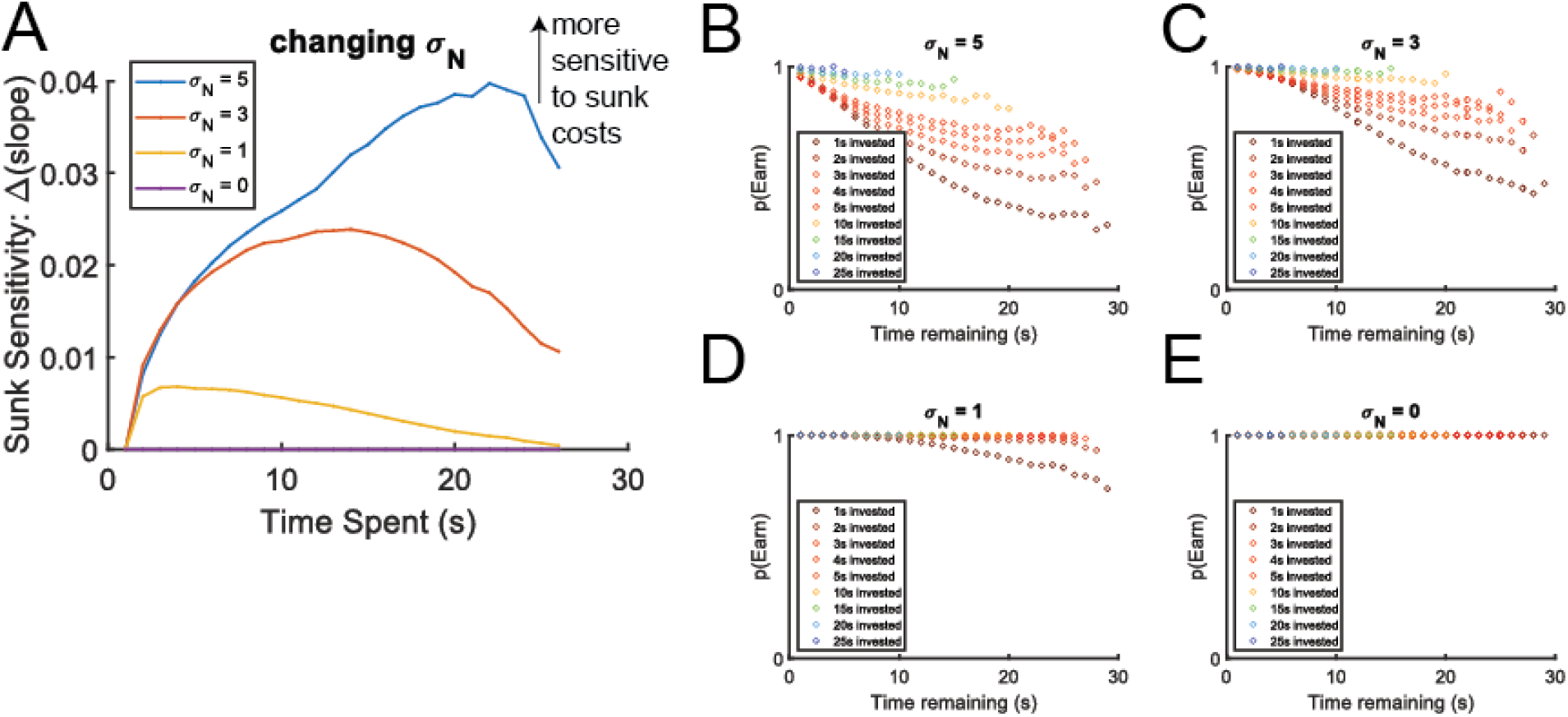
Changing σ_N_. As we change the speed at which the willingness to wait ***W*** drifts, the sensitivity to sunk costs changes, but in ways never observed in real data. (A) The sensitivity to sunk costs measured as the difference between the slopes as measured for a given time-spent and the control analysis measuring the slope of the 0-time-spent condition of the same time-remaining points that go into the slope measurement for a given time-spent. This equals the difference between the blue stars and black circles, respectively, shown in Figure 4B,D. If there were no sensitivity to sunk-costs, this difference would be 0. Sunk costs are indicated by a positivity in this measurement (observed slope > control slope), reflecting the amount of increase in ***pEarn*** due to sunk costs (escalation of commitment in the wait phase). (B-E) Slope panels showing the effect of changing σ_N_. The original formulation has σ_N_=3. Note that when σ_N_=0, there is no sensitivity to sunk costs because the agent never quits. **<u>*Results shown are from our simulations using the Ott et al. model*</u>**.

**FIGURE 8:**
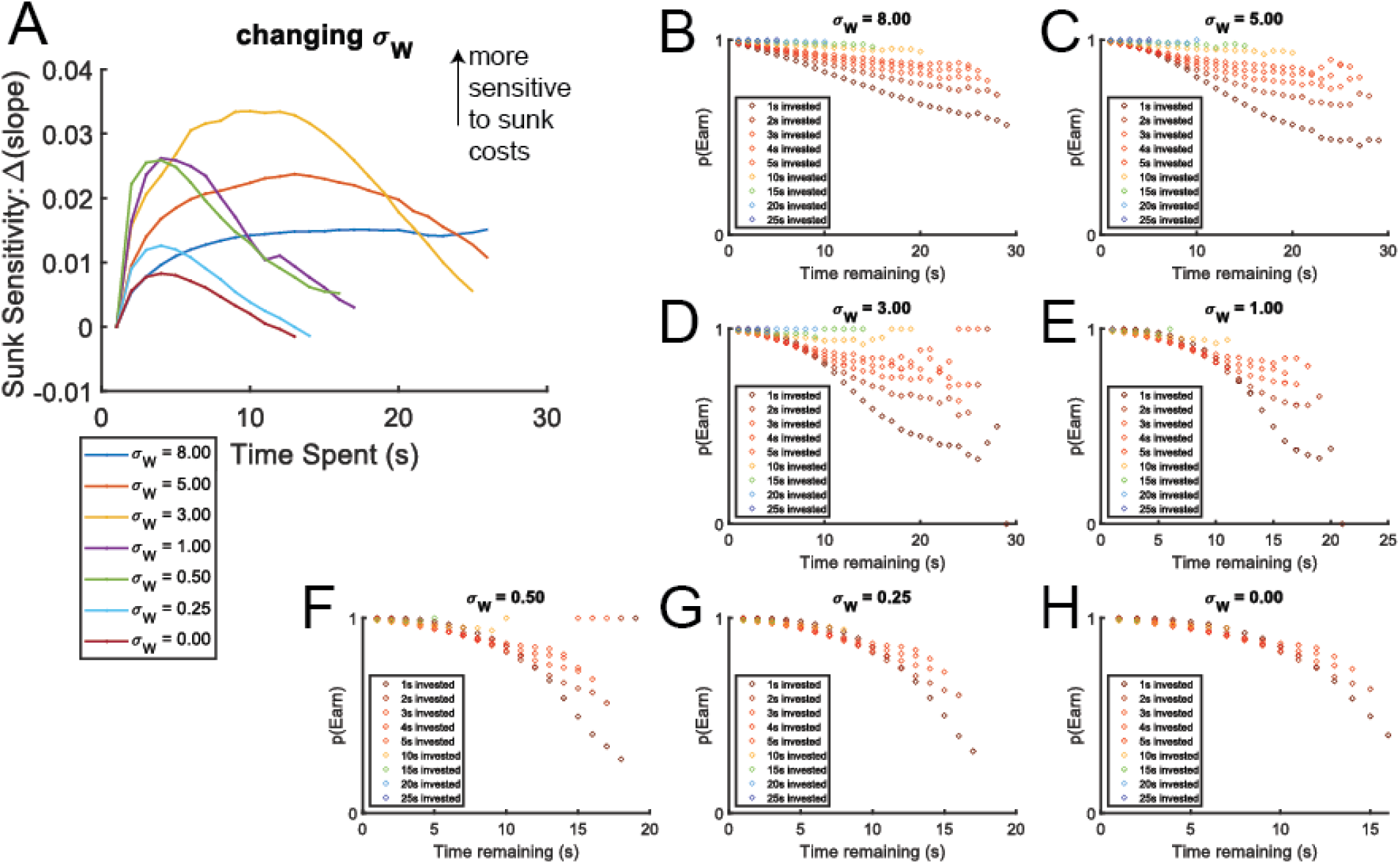
Changing σ_W_. As we change the variation of the ***W*** parameter around the threshold, the sensitivity to sunk costs changes, but in ways never observed in real data. (A) The sensitivity to sunk costs measured as the difference between the slopes as measured for a given time-spent and the control analysis measuring the slope of the 0-time-spent condition of the same time-remaining points that go into the slope measurement for a given time-spent. This equals the difference between the blue stars and black circles, respectively, shown in Figure 4D. If there were no sensitivity to sunk-costs, this difference would be 0. Sunk costs are indicated by a positivity in this measurement (observed slope > control slope), reflecting the amount of increase in ***pEarn*** due to sunk costs (escalation of commitment in the wait phase). (B-E) pEarn panels showing the effect of changing σ_N_. Compare Figure 4A. The original formulation has σ_W_=5. Note that when σ_W_ is very small, the sensitivity breaks down as there is little variation for the slope calculations to latch onto. **<u>*Results shown are from our simulations using the Ott et al. model*</u>**.

However, as can be seen in Figure 8, changing σ_W_ does not provide any reason for the delayed accrual of sunk costs seen in the RD task. In contrast, our dual-decision explanation provides a parsimonious explanation for both the changes in sunk costs and the delayed accrual.

We tried modeling the delay to start in two ways. First, we tried modeling it by setting σ_N_ to 0 for the first few seconds. Second, we tried modeling it by setting the quit-threshold TWZ to 0 for the first few seconds, but had the model accept every offer (see Figure 12).

**FIGURE 9:**
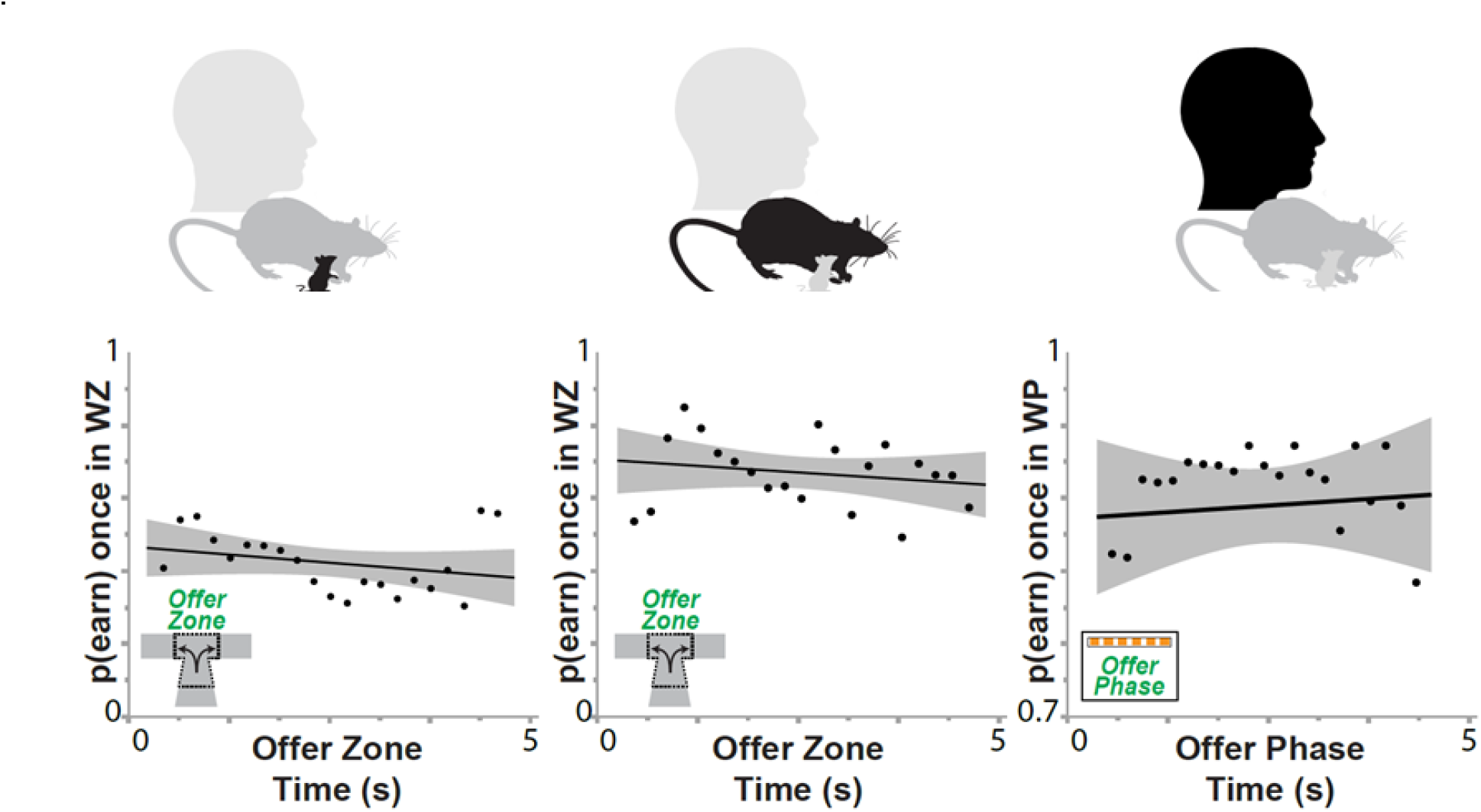
No sunk costs were seen in the offer zone. The time spent in the offer zone had no effect on the probability of waiting out the delay once the subjects reached the wait zone. (Left: mice; center: rats; right: humans.) Figure from (Sweis, Abram, et al. 2018). Used with permission of the publisher.

**FIGURE 10:**
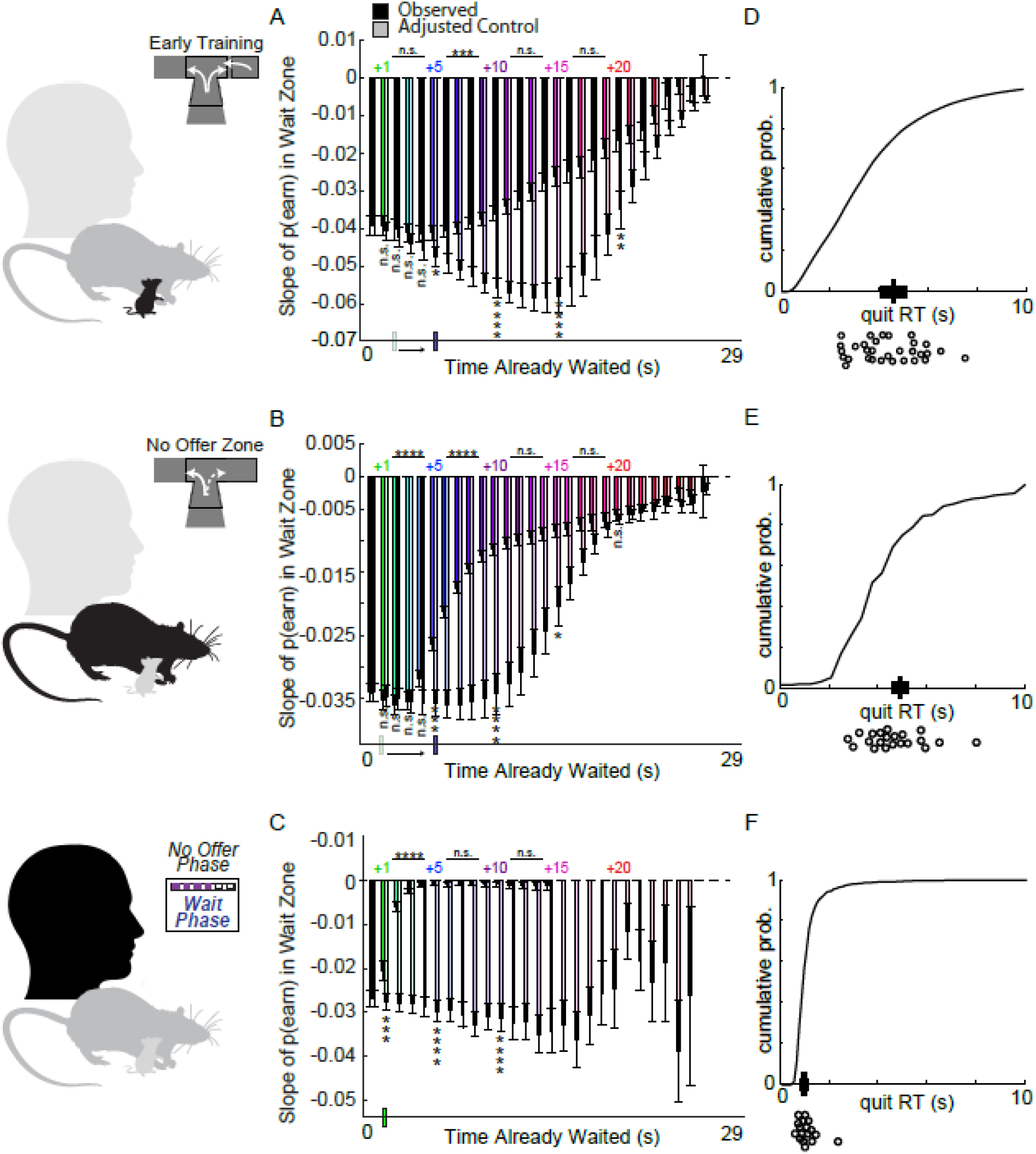
In variants with no offer zone, the start of the sunk cost accumulation is delayed at a similar time-frame to the typical quit time. (Remember, because there is no offer zone in these variants, all choices are quit vs earn. It is our contention that these early quits are actually skips.) (A,B,C) Colored bars show the slope of **p(Earn)** for each value of Time spent, open bars show the equivalent control. (D,E,F) The distribution of quit times peaks at the moment that sunk cost sensitivity starts to appear. Figure from (Sweis, Abram, et al. 2018). Used with permission of the publisher.

**FIGURE 11:**
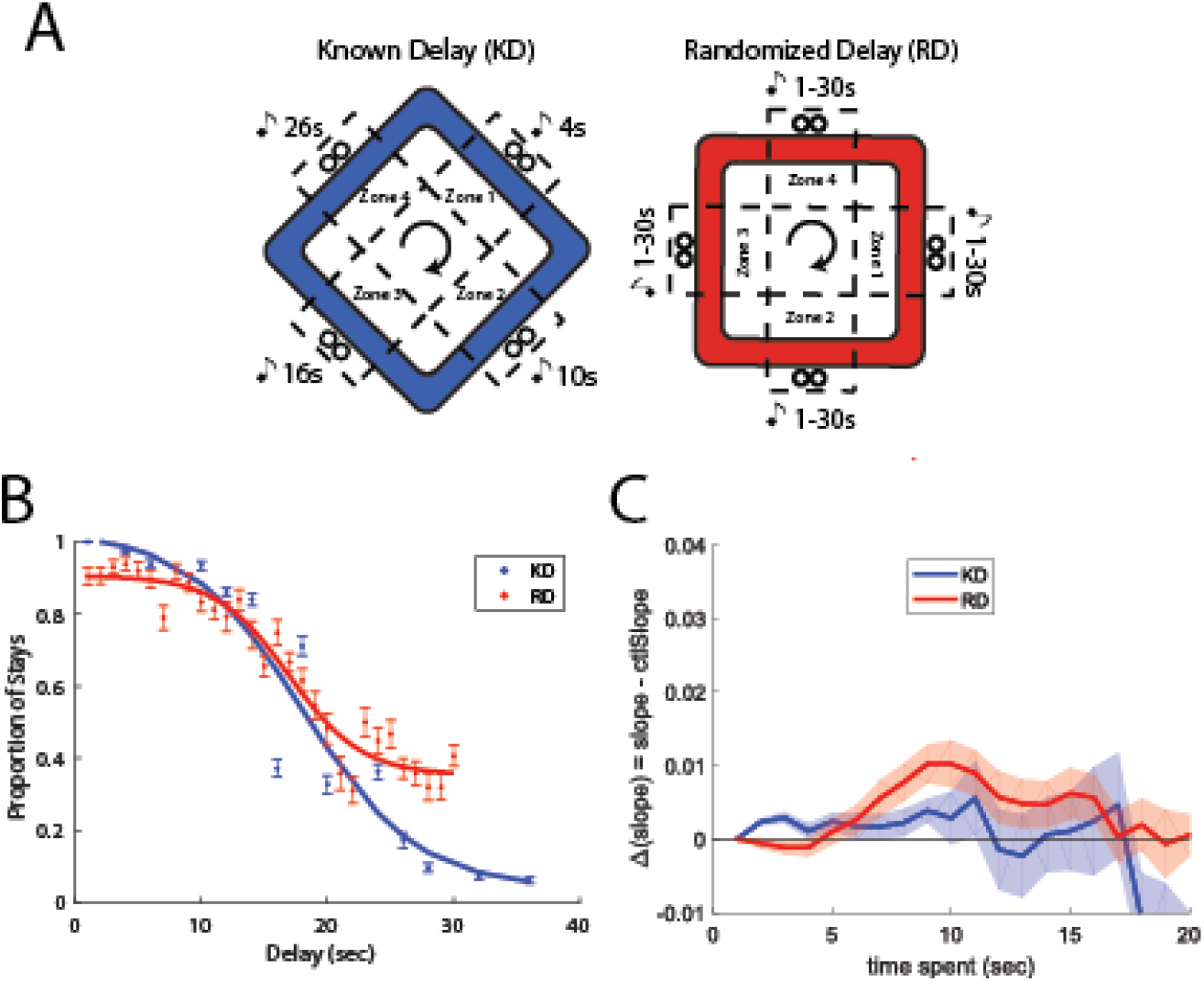
Sunk cost sensitivity in the Known-Delay / Randomized-Delay tasks. (A) The two tasks. In these tasks, rats encountered four restaurants providing the same flavor of food. There was no separate offer zone, but rats could leave the zone before food was delivered, rescinding the offer. In the Known Delay condition, the delays were constant within a restaurant each day, but changed between restaurants and between days. In the Random Delay condition, the delays were random from 1s-30s. Rats did not know the delay in the Random Delay condition until entering the restaurant. (B) Rats showed more consistency with a sharper response curve on the Known Delay task than on the Random Delay task. (C) Sunk cost sensitivity was larger on the Random Delay condition than on the Known Delay condition. Sunk cost sensitivity also was delayed in the Random Delay condition relative to the Known Delay condition, suggesting that there was a separate decision time, as seen in the Restaurant Row conditions without an offer zone. From (Duin et al. 2021). Used with permission of the publisher.

**FIGURE 12:**
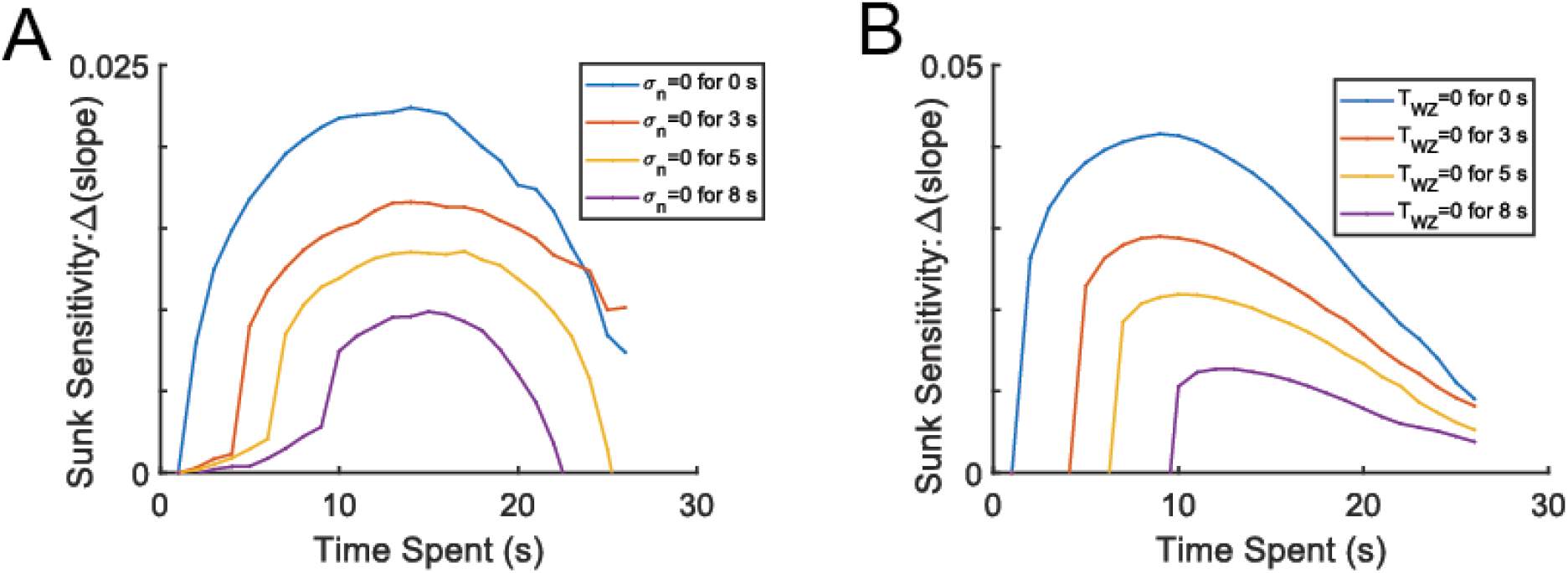
Modeling the delay before sunk costs start accruing in the wait zone. (A) Setting σ_N_ to 0 for the first few seconds prevents the agent from quitting because the agent only takes offers with ***W***_**0**_ > offer, so the animal won’t quit. (B) Forcing ***T***_WZ_ to 0 and taking every offer puts a hard inability to quit while ***T***_WZ_ is 0. Both models produced less sunk costs, the opposite of what was seen in (Duin et al. 2021). Compare to Figure 11. **<u>*Results shown are from our simulations using the Ott et al. model*</u>**.

If we model the delayed start explicitly in the Ott et al. framework, we find that it decreases the sunk costs, which is the opposite result of that seen in the KD/RD comparison. We modeled this delayed accrual start by setting σ_N_ to 0 for the first few seconds, which effectively made the animal extremely unlikely to quit (as ***W***_**0**_ was known to be greater than the offer by definition - that’s why the animal accepted the deal in the first place). This model showed much smaller sunk cost effects after the delay.

If we model the delayed start by directly setting the quit-threshold ***T***_WZ_ to 0 for the first few seconds while having the model accept every offer, we found that this reduced the sunk costs after the delay. Accepting every offer removed the offer phase from the model. Setting the quit-threshold to 0 for the first few seconds made the model not start accruing sunk costs until the delay had ended. This dramatically reduced the sunk costs seen after the delay, again, incompatible with the KD/RD results.

### 3. The data finds a lack of sensitivity to sunk costs in some conditions

The Ott et al. model argues that the sunk cost sensitivity observed in the Restaurant Row and Web Surf tasks are due to fundamental aspects of the task construction (the attrition effect due to the interaction of the sampled willingness-to-wait ***W*** against the offer phase threshold ***T***_OZ_ and the wandering ***W*** against the changing quit threshold ***T***_WZ_). This means that every agent should show sunk costs in the Restaurant Row and Web Surf tasks. However, we have found several conditions where agents (mice, rats, humans) do not show a sensitivity to sunk costs, which is, on the surface, incompatible with the Ott et al. model.

When humans were distracted from the delay countdown, we found that the sunk costs disappeared (Kazinka, MacDonald, and Redish 2021). Similarly, in rich environments (when the offer distribution is such that rodents can easily earn a large amount of food within the environment), we do not see sunk costs (Sweis, Thomas, and Redish 2018). In these conditions, we find that subjects show identical slopes between conditions, independent of the time spent waiting. These data imply a lack of sensitivity to sunk costs, even though the animals do continue to quit. Thus, quitting behavior can occur in the absence of sunk costs.

While these situations can be modeled with a constant, non-zero probability of quitting, we have not found any reasonable change in the Ott et al. model that can produce this constant probability of quitting. The most obvious change to make, removing the rate of diffusion of **W** through time (decreasing σ_N_), does not produce this effect. As can be seen in Figure 7, It does reduce sunk costs, but by driving all decisions to waiting out the delay, which is inconsistent with the observed data in the distracted and reward-rich conditions.

The Ott et al. model does include thresholds, value, and offer-decisions, so we could directly model the 1s-15s reward-rich world in which mice accept all offers in the offer zone. In real data, mice do not show sunk costs in this condition. However, the Ott et al. model continues to show robust sunk costs that are no different from the 1s-30s economically scarce condition.

The sunk cost effect is occurring in their model because the subjects are not accepting all of the 1-15s offers in the offer zone. With a ***W*** centered at 18s and a non-zero σ_W_, a few of the offers are not being selected. However, even if subjects are taking all of the offers, we still see sunk costs in their model.

The only way we were able to construct simulations that showed similar quitting phenomena to our distracted humans (or mice in rich environments) was by changing the slope of the quit-threshold function to a negative number, implying that there was an <u>increased</u> likelihood of quitting the longer the subject remained within the wait phase.

Presumably the result in Figure 16 arises because, in the Ott et al. model, sunk costs depend on both the attrition of the set of offers accepted in the offer zone and from the sloped quit threshold in the wait zone. Thus, setting the quit threshold to increase quitting probability with time (anti-sunk-costs) can counteract the effect of attrition in the set of offers accepted on sunk costs in their model. We find this hypothesis to be unreasonable. Presumably, Ott et al. could argue that distracted subjects or subjects living in a rich environment are making *quit* decisions via different decision processes (as we have argued for the offer phase), but we find this to be an implausible assumption, and argue instead for the more parsimonious hypothesis that sunk costs depend on attention to the task at hand because it is about recognizing the decaying delay.

**FIGURE 13:**
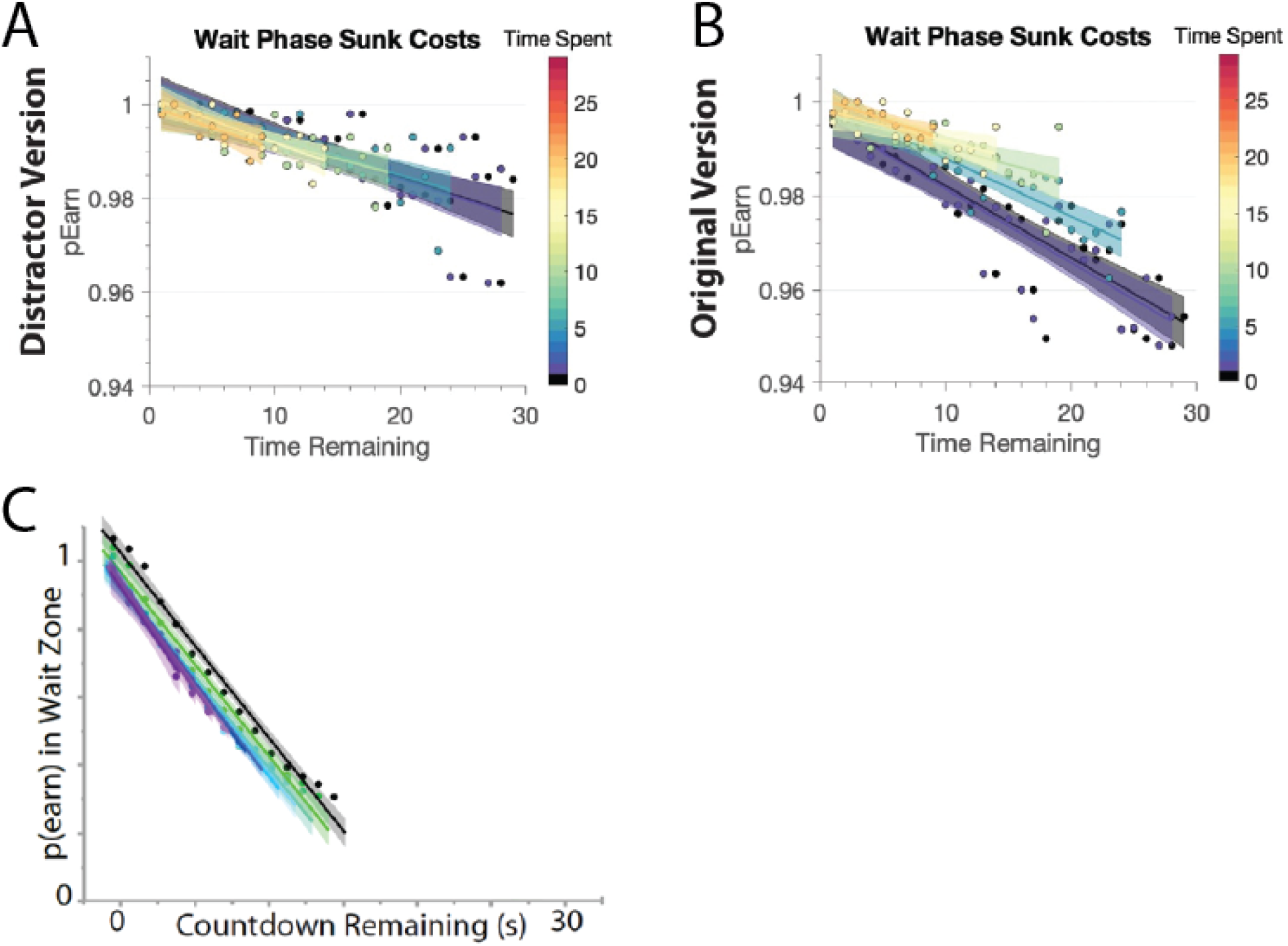
Quitting behavior can be insensitive to the sunk costs. Neither the Restaurant Row nor the Web Surf tasks automatically produce a sensitivity to the sunk costs. (A) Quitting behavior in humans in the distractor version of the Web Surf task in an online sample from (Kazinka, MacDonald, and Redish 2021). Original article distributed under CC-BY. Compare (B) quitting behavior in a replication of the original version from another online sample. (C) In a resource-rich version of the Restaurant Row (during training, when delays only ranged from 1-15s), mice also showed quitting insensitive to the sunk costs. From data reported in (Sweis, Abram, et al. 2018).

**FIGURE 14:**
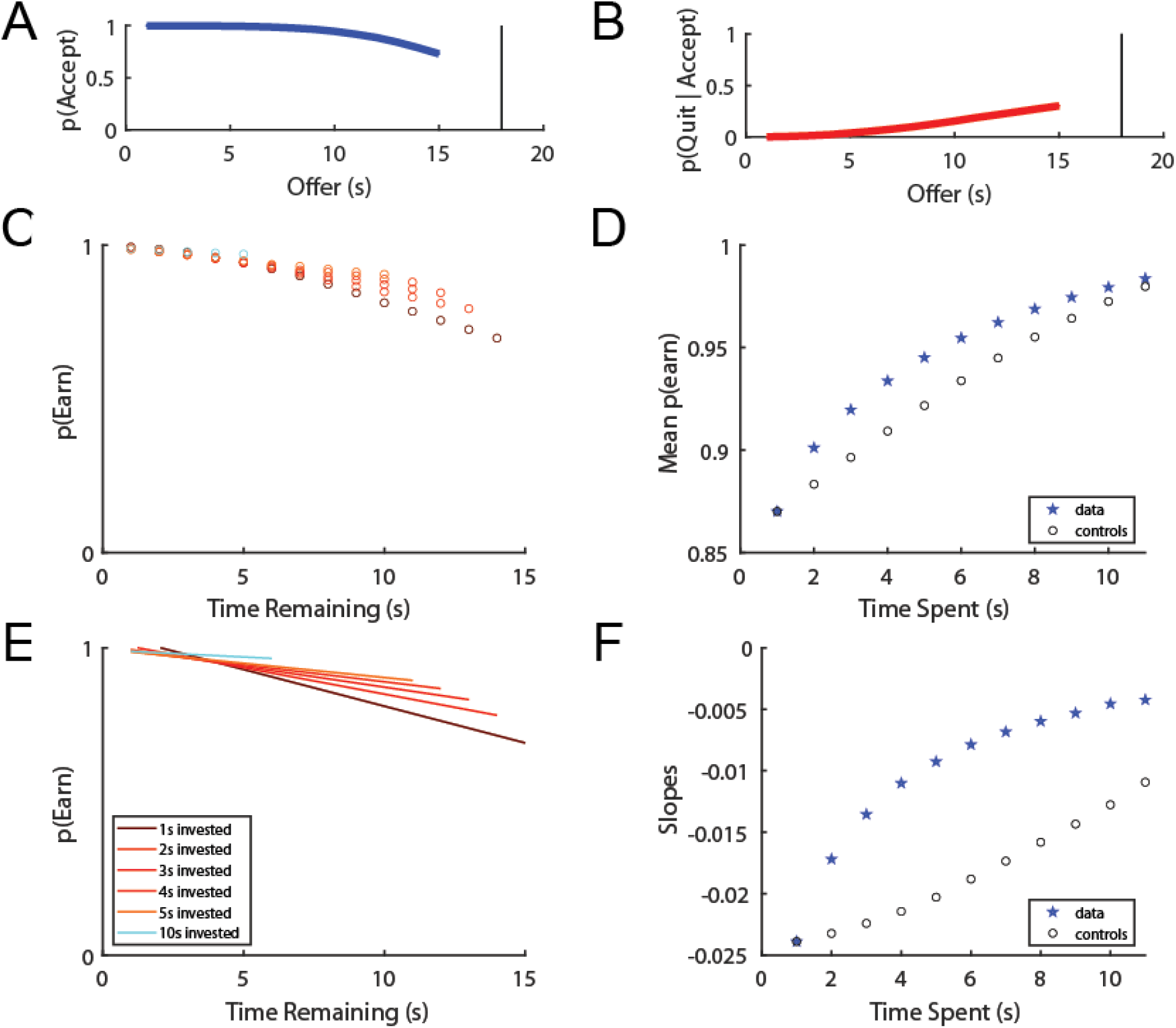
Simulation of the Ott et al. model when offers only range from 1s-15s. Note the continued presence of a sensitivity to sunk costs. **<u>*Results shown sare from our simulations using the Ott et al. model*</u>**.

**FIGURE 15:**
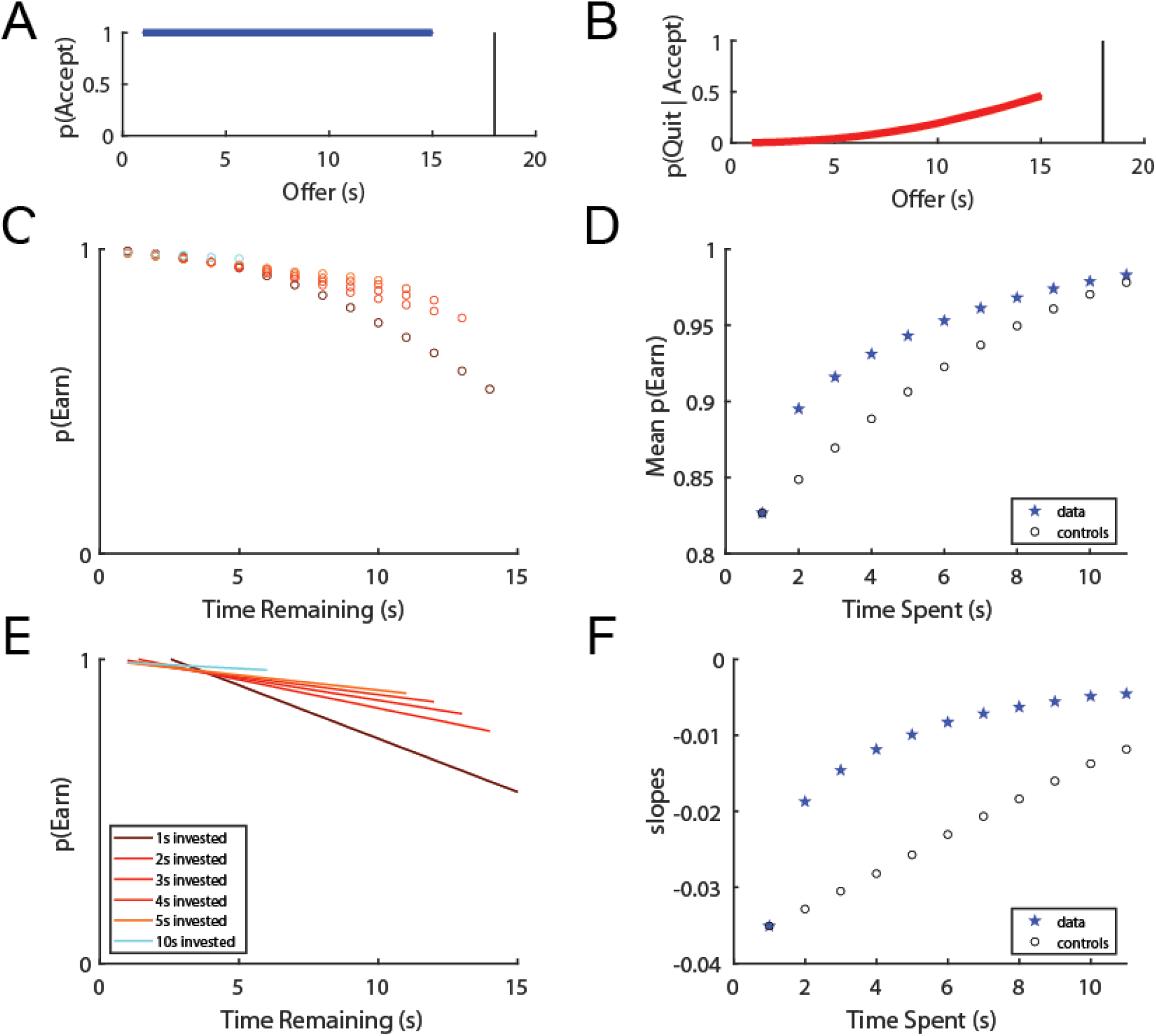
Simulation of the Ott et al. model when offers only range from 1s-15s, and the agent accepts all offers in the offer zone. Note the continued presence of a sensitivity to sunk costs. <u>***Results shown are from our simulations using the Ott et al. model***</u>.

**FIGURE 16:**
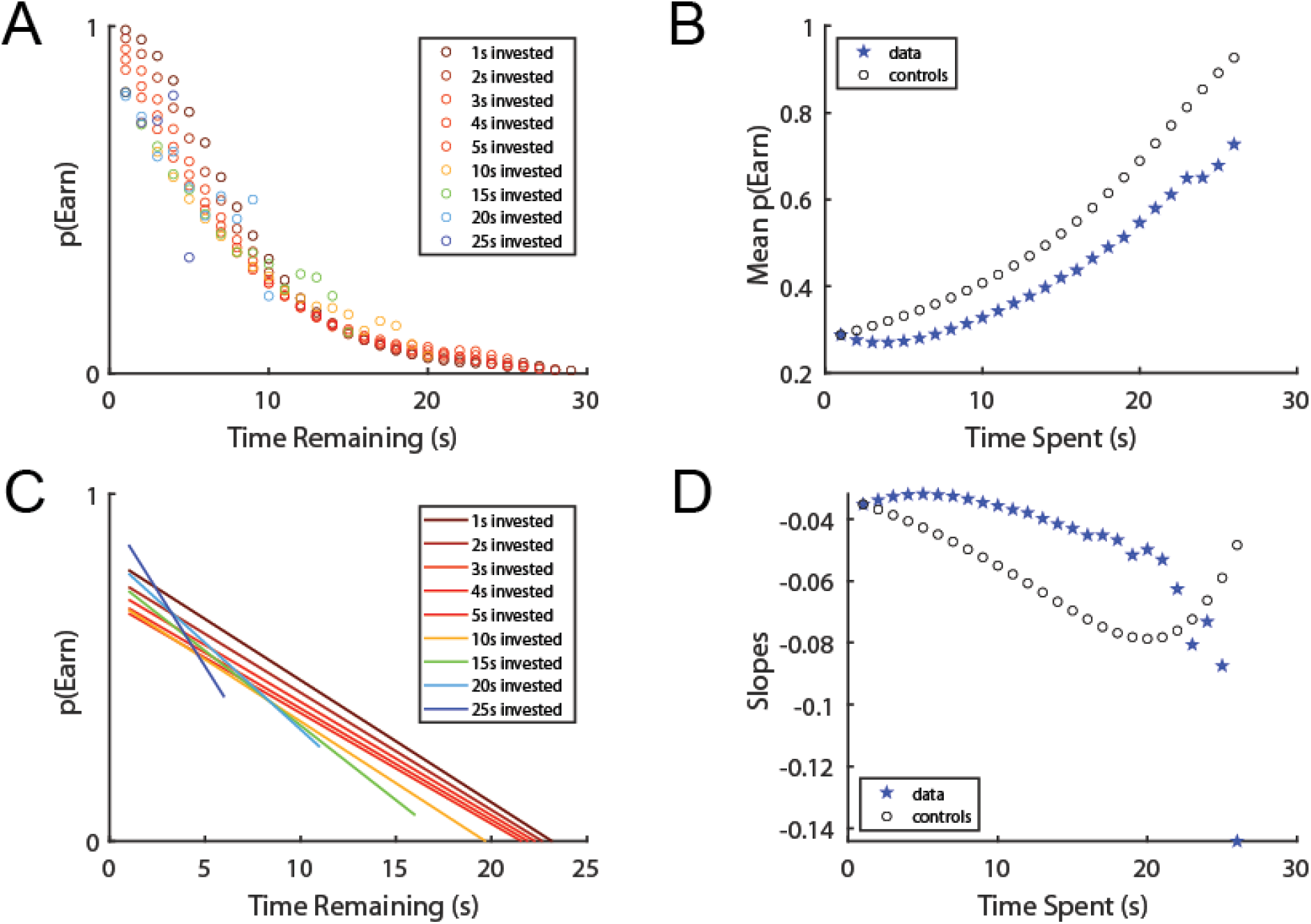
The only way we were able to construct a probability of quitting that was essentially constant over the time invested was to change the slope of the quitting threshold to be <0. This makes the threshold RISE over time, and is, we believe, a nonsensical variant of the model. **<u>*Results shown are from our simulations using the Ott et al. model*</u>**.

Importantly, however, the wait phase thresholds (*stay*/*quit*) do not generally change in mice between the 1-15s and 1-30s phases (Sweis, Thomas, and Redish 2018), implying that it is unlikely that they are making decisions through vastly different mechanisms in these conditions. Nevertheless, we do see sunk costs arise in the 1-30s situations but not in the 1-15s situations. Given that the Ott et al. model depends on the relationship between willingness to wait and quit-thresholds in the wait zone, their model would predict similar sunk costs in both situations. An alternate explanation is that the sunk costs are coming from subjects having difficulty leaving a bad offer that they decided to take.

### 4. The basic model does not fit the structure of the raw data

We do not observe sunk costs in all conditions. Nor do we observe sunk costs at all times within the waiting phase. Nor do we observe sunk costs uniformly within the waiting phase. Instead, what we have observed is that the sensitivity to sunk costs primarily occurs when subjects take bad deals, before the deal transitions to becoming a good deal, suggesting that quitting is a correction of a mistaken choice made in the offer phase.

A good offer is one that is below one’s threshold for that flavor / gallery. (The offer is of lower cost than the agent is typically willing to spend for that reward.) A bad offer is one that is above one’s threshold for that flavor / gallery. (The offer is of higher cost than the agent is typically willing to spend for that reward.) In practice, we have observed that quits tend to occur in situations where the agent accepted a bad (high-cost, above-threshold) offer. This means that quits are typically correcting mistakes made in the offer zone (Sweis, Abram, et al. 2018; Sweis 2018).

In Figure 17, we rearranged the sunk cost measure by the value that was left at the countdown at the time of quitting. (Value is defined as the difference between the threshold that that agent has for that flavor / gallery and a given offer: V = threshold - offer. Thus high offers have low value and low offers have high value.) As can be seen, the probability of waiting out the delay to earn is enhanced through sunk costs **only** in the bad deals (deals where the delay remaining was above the typical threshold for that reward site [restaurant or gallery]). These data show that a sensitivity to sunk costs comes under conflicting decisions of whether to wait out an excessive delay that was accepted incorrectly or to quit, thereby wasting the time already spent. Once the delay during the countdown has crossed the typical threshold, the deal is now seen as reasonable, even if the already spent time is lost, and the sensitivity to sunk costs disappears. We argue that the conflict of whether to rectify the incorrectly accepted deal is driving the sensitivity to sunk costs because the wasted costs lost by rectifying the incorrectly accepted deal are the time already spent (the sunk costs).

**FIGURE 17:**
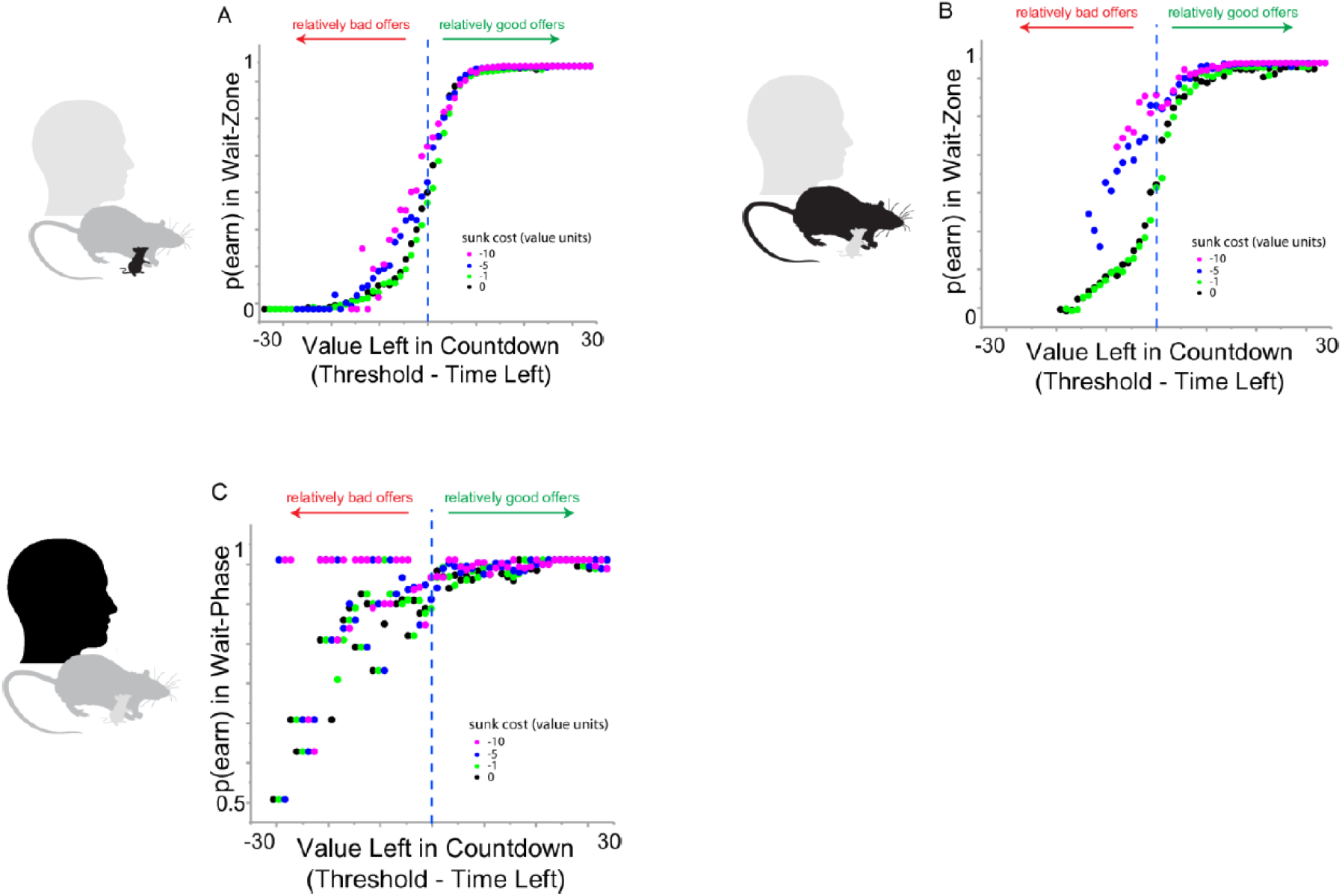
Sensitivity to sunk costs depends on the value remaining. Value = threshold - offer. The x axis shows the value left in the countdown at the time of quit. The colored dots show the time already spent, converted into value. Colored dots indicate the time spent, Pink = -10s, blue = -5s, green = -1s, black = 0s. Thus pink dots have more sunk costs spent than blue dots, which have more than green. Black dots indicate no sunk costs yet spent. The vertical line marks the transition to where the value crosses zero, that is where the delay crosses threshold. Thus to the right of the line, the remaining time is below the threshold for that restaurant / gallery. Mice, rats, and humans were only sensitive to sunk costs until the value crossed that line. From (Sweis 2018).

**FIGURE 18:**
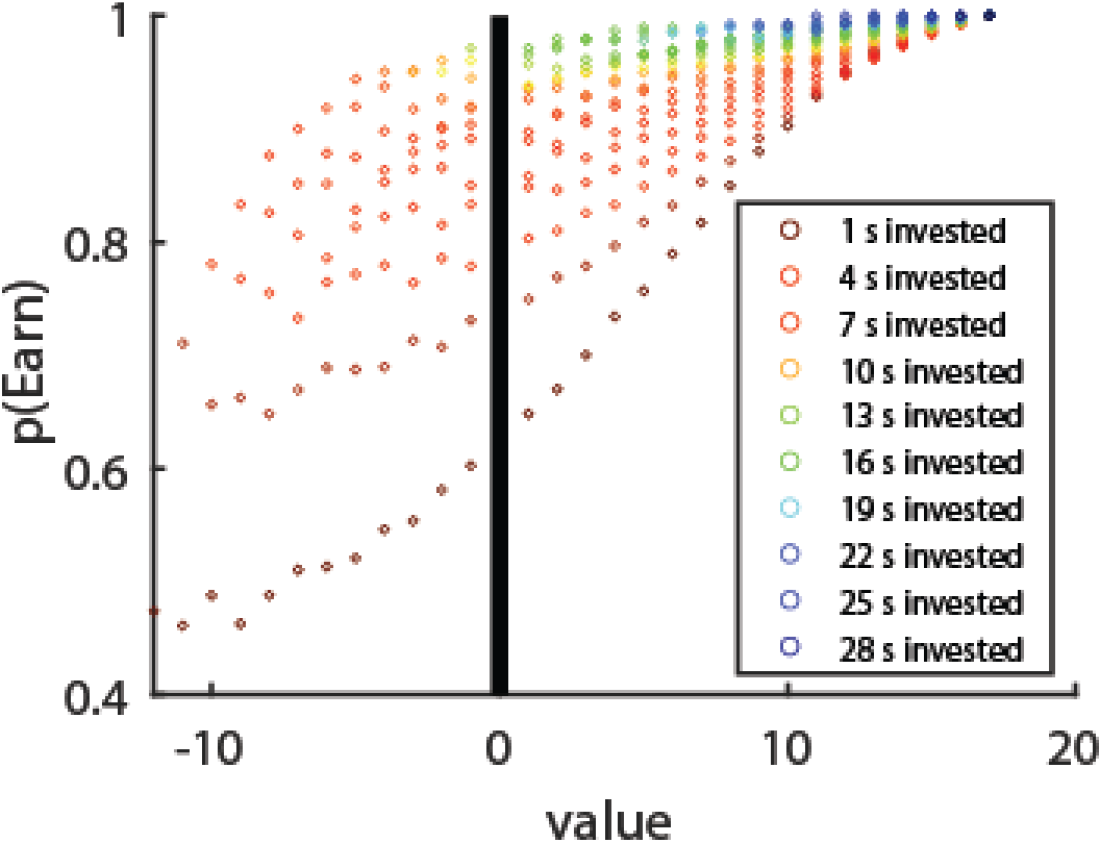
Replotting the original Ott et al. simulation (see Figure 4) in the format used in FIGURE 17. Note that the simulation is unaware of the switch between negative to positive value and thus continues showing sunk costs across the whole time, which is not what is seen. **<u>*Results shown are from our simulations using the Ott et al. model*</u>**.

Because the Ott et al. model is uniform, there is no way for it to model these results. Their model shows a uniform exponential fall-off of quitting, independent of the future outcomes. It does not differentiate rectifying having accepted a bad deal before crossing the threshold from rectifying it after crossing that threshold.

## Discussion

As can be seen from the data analyses above, the Ott et al. model (Ott et al. 2021) cannot explain the behaviors we see within variants of the Restaurant Row and WebSurf tasks. We therefore conclude that the sunk cost sensitivity is a real effect showing a sensitivity to sunk costs (as defined by an escalation of commitment, (Staw 1976; Staw and Fox 1977; Staw and Ross 1989)) and is not an epiphenomenal consequence of the simple explanation suggested by Ott et al. (Ott et al. 2021).

While we commend Ott et al. for the simplicity of their model, it cannot account for the behavioral phenomena observed in our tasks. As noted, they are able to show sunk cost-like effects with a simple model. However, theirs is not the first simple model to show a sensitivity to sunk costs. In fact, several previous animal learning theory models have been put forward that produce sunk cost effects, including the state-dependent-valuation learning [SDVL, (Kacelnik and Marsh 2002; Pompilio, Kacelnik, and Behmer 2006; Aw, Vasconcelos, and Kacelnik 2011)] and within-trial-contrast [WTC, (Singer and Zentall 2011)] models. The SDVL model notes that ongoing energy expenditures imply that food rewards will become more valuable to an agent as time progresses. Thus, reward valuation should depend on the time since the last time reward was received. Of course, the SDVL model cannot explain why humans foraging for videos show similar effects, but we can hypothesize a more general “value process” that depends on recency of experience. The WTC model notes that distant rewards are discounted so that approaching rewards are seen as increasing in value, which can provide an increasing contrast between the current state of the animal (changing through SDVL) and the approaching reward (Singer and Zentall 2011). In fact, the Ott et al. model contains components of the SDVL and WTC models within it in the quit-threshold ***T***_WZ_ that slopes down from the threshold at entry to zero at the time of reward. As shown above, a part of the sunk cost effect seen in the Ott et al. model is due to the slope of this quit threshold. Of course, as noted above, none of these models can explain the lack of effects in the offer phase.

We suspect that a more parsimonious explanation of the data is the increased motivation provided by Pavlovian associations between the countdown in the wait zone and the value of the outcome. This would show decreases with decreased attention (Kazinka, MacDonald, and Redish 2021), as well as sensitivity to factors that increase that relationship to the reward zone (such as conditioned place preference) (Robinson and Berridge 1993; Berridge 1996; Clark, Hollon, and Phillips 2012; Fanselow and Wassum 2015). This is a form of the endowment effect (Kahneman, Knetsch, and Thaler 1990, 1991; Redish, Schultheiss, and Carter 2016) and is similar to the deliberative vs implementational mindset constructs in the human literature (Gollwitzer, Heckhausen, and Steller 1990; Achtziger and Gollwitzer 2007). Essentially, this hypothesis suggests that quitting out of the wait zone is a recognition of a mistake and staying is due to an unwillingness to leave that mistake. Because the extent of the mistake depends on the effort spent (more effort spent was a larger mistake), the decision becomes related to the effort already spent and shows a sensitivity to the sunk costs.

It is worth noting that there are situations where an increased motivation from effort already spent can actually be economically useful (see (Inzlicht, Shenhav, and Olivola 2017) for discussion), which could further provide evolutionary drive to make decisions based on past costs. There are many cases where sunk costs can provide the additional incentive to push through difficult choices. For example, when pushing through the last part of a marathon run, a common coaching line is “you’ve come so far already, you can finish!” Similar incentives can be seen in the last segment of any long task that requires pushing through burnout and exhaustion, such as a PhD thesis. In truth, the correct comparison is between quitting (with no reward but no additional costs) and continuing (with the reward of finishing but with the additional costs of pushing through the burnout). It is possible that the costs of pushing through may appear over-burdensome because of economic myopia. The additional motivation provided by the refusal to quit (due to sunk costs) can allow one to achieve goals that may be difficult to achieve otherwise. This motivational effect can be interpreted as the following logic: in a long marathon, there is uncertainty in that future valued goal. Thus it becomes easy to conclude that the goal is not worth the additional effort, particularly as that additional effort increases with exhaustion. But because the sunk costs provide additional evidence that the goal will be worth it, the past effort spent provides increased motivation.

In this response, we have pointed out aspects of the observed data that are incompatible with this simple model. We conclude that the temporal sunk costs observed in these tasks is not an epiphenomenon of simple changes in state (SDVL, WTC, the quit-threshold slope in Ott et al.) nor is it an epiphenomenon consequence of attrition effects due to the observed offer thresholds. Instead, these data imply that the sensitivity to temporal sunk costs seen in these tasks is an example of escalation of commitment akin to that studied in the business and human behavior literatures (Staw 1976; Staw and Fox 1977; Staw and Ross 1989).

## Methods

Simulations were based on those of <u>Ott et al. (2021)</u>. A simulated agent was designed that made two decisions: first an enter/skip decision (modeling the offer zone / offer phase in Restaurant Row and WebSurf) and then a repeated quit/remain decision (modeling the wait zone / wait phase). In a given trial, the agent started with a draw of a willingness to wait, **W**, drawn from a normal distribution around the threshold (**W**_**0**_=**T**_**OZ**_ defined as 18s) with standard deviation **σ**_**W**_: W = W_0_ + N(σ_W_). If **W** was greater than the threshold (W>T_OZ_), the agent accepted the offer and “entered the wait zone”. In entering the wait zone, the willingness to wait W drifted each second by adding in normally distributed random noise with standard deviation **σ**_**N**_. A quit-threshold was defined as the linear function such that **T**_**WZ**_=**W**_**0**_ at the start of the waiting time and decreasing by 1s each second until it reached **T**_**WZ**_=0 at the end of the waiting countdown. If the cumulative willingness to wait (integrating the drift over time) ever fell below the quit-threshold TWZ, the agent “quit” out of the wait zone. Trials were independent.

Initial parameter set tested: **W**_**0**_ = **T**_**OZ**_ = 18s. **σ**_**W**_ = 5s. **σ**_**N**_.= 3s. Offers were uniformly distributed over 1s-30s inclusive.

Variations of these parameters were run as follows.

Figure 4: Original model as described above.

Figure 5: Instead of accepting if **W**>**T**_**OZ**_, all agents always accepted all of the offers.

Figure 6a/b: Original model as described above.

Figure 6c/d: Instead of sloping down from **T**_**OZ**_ to 0, the quit-threshold was set at a constant **T**_**WZ**_=**T**_**OZ**_.

Figure 6e/f: Instead of sloping down from **T**_**OZ**_ to 0, the quit-threshold was set at a constant **T**_**WZ**_=0.

Figure 7: Tested multiple potential values for **σ**_**N**_: 0s, 1s, 3s, 5s.

Figure 8: Tested multiple potential values for **σ**_**W**_: 0s, 0.25s, 0.5s, 1s, 3s, 5s, 8s.

Figure 12a: **σ**_**N**_ was set to 0 for the first k seconds. Because the agent only accepts the initial offer if **W** > **W**_**0**_ = **T**_**OZ**_ = **T**_**WZ**_(start), **W** > **T**_**WZ**_ for those first k seconds, and the agent will not quit for those first k seconds. Then W starts drifting. However, it starts from the W that it was when the agent entered the wait zone, even though **T**_**WZ**_ has already decreased by k.

Figure 12b: **T**_**WZ**_ was set to 0 for the first k seconds. Because **W** >> 0 at entry and **σ**_**N**_ is small, the agent will not quit for those first k seconds. After k seconds, **T**_**WZ**_ was returned to where it would have been after k seconds, but **W** has drifted for those k seconds as in the original model.

Figure 14: Offers were uniformly distributed over 1s-15s inclusive. Note that although **W**_**0**_ = 18s > 15s, because **W** + N(**σ**_**W**_) could be < 15s.

Figure 15: Offers were uniformly distributed over 1s-15s inclusive and agents accepted all offers.

Figure 16: Instead of **T**_**WZ**_ decreasing by 1s per second, **T**_**WZ**_ increased by 1s per second.

Figure 17: Original model as described above.

1 million trials were run for each experiment. Simulation code is available at https://github.com/adredish/SunkCostModels-Redish2021.

